# Enhanced Complex Stability and Optimal JAK Geometry are Pivotal for a Potent Type III Interferon Response

**DOI:** 10.1101/2022.09.27.508945

**Authors:** Theint Aung, Curt M. Horvath, Juan L. Mendoza

## Abstract

Type I and III Interferons (IFN) constitute the host system’s first line of defense against viral infections. Although the two families use distinct extracellular receptor complexes, an identical pair of Janus kinases (JAK) activates a similar set of signal transducer and activator of transcription (STATs) through a conserved pathway. Consequently, type I and III IFNs are presumed to activate a largely overlapping set of IFN-stimulated genes (ISGs) and elicit similar biological responses. Therapeutically, type III IFNs are attractive alternatives to type I IFNs due to their innate tissue specificity and lower systemic toxicity. However, a major limitation of type III IFNs is the significantly lower potency of their physiological activities compared to type I IFNs. To evaluate the role of receptor geometry in IFN signaling, we engineered cell lines that express wild-type and mutant IFNλ receptors (IFNλR1) with rotated intracellular registers with respect to the associated JAK1. Evaluation of downstream signaling and biological activity of type III IFNs in cells with varied JAK-JAK geometries uncovered variant receptors that result in potentiation of type III IFN signaling and all downstream activities. With the combined use of a high-affinity ligand to stabilize the extracellular receptor complex, we found that the optimization of the intracellular JAK-JAK geometry enhances the type III IFN antiviral and anti-proliferative activities by 2- and 3-logs, respectively. In addition to providing a molecular basis for the observed differences in potency between type I and III IFNs, this work provides deeper insights into broader cytokine signaling mechanisms and novel blueprints for modulating cytokine functions to develop next generation antiviral and anticancer therapeutics.

## Introduction

Type I and III Interferons (IFNs) are two distinct cytokine families that are crucial in arming the host system with an efficient and controlled state of immunity in response to infections^1,2^. Both IFN families share important biological functions^3,4^ that modulate the innate and adaptive arms of the immune system to activate gene expression programs involved in antiviral, anti-proliferative, anti-tumoral and other immunomodulatory pathways^5,6^. For both families, the cytokine production is similarly induced by cellular sensing of pathogen-associated molecular patterns (PAMPs) from viral or non-viral pathogens via pattern recognition receptors (PRRs)^7,8^. The upregulation of type I and III IFNs expression then initiates a chain of signaling cascades that lead to robust transcriptional induction of related IFN-stimulated genes (ISGs)^9,10^. Although there is a significant overlap in the pool of ISGs induced between type I and III IFNs, multiple studies have shown that the two pathways are non-redundant, but rather have complementary functions that serve to maintain an optimal state of immune protection^11–13^.

To understand and address the discrepancies in activity between the type I and type III IFNs, our previous studies on type III IFNs focused on the affinity of the ligand to its receptors composed of IFNλR1 and IL10Rβ. Through directed evolution, we engineered a high-affinity variant (IFNλ3 H11) that we tested for the ability to potentiate activity^14^. These studies revealed that the affinity of the receptor complex, while a contributing factor, only partly explained the lower activity of type III IFNs. More importantly, we found that the signaling amplitude (E_max_) elicited by the type I IFNs is three times greater than the signal amplitude elicited by the type III IFNs for which the extracellular receptor affinity had no effect. We hypothesized that this difference in signaling amplitude likely explains the weak IFN functions such as ISG expression, antiviral and anti-proliferative/anti-cancer activities.

Gene expression studies have shown that type III IFNs induce a weaker transcriptional profile of a smaller set of ISGs compared to type I IFNs^15–17^. Consequently, type III IFNs are less efficacious as antiviral or anti-tumoral agents compared to type I IFNs^18–20^. Such a significant gap in potency currently presents an insurmountable hindrance in translating type III IFNs for clinical uses^21^. This is reflected by the fact that no type III IFNs has been approved for clinical use whereas many type I IFNs are already clinically utilized to treat cancers, autoimmune disorders, and viral infections^22–24^. However, the therapeutic gains of type I IFNs are unfortunately offset by the adverse side effects in patients^25^. Given the much more favorable toxicity profile of type III IFNs largely due to the restricted expression of type III IFN receptors to epithelial and barrier cells, there is much vested research interest in understanding the key factors contributing to the lower potency of type III IFNs so that new strategies can be developed to overcome these limitations^26^.

Notably, the differential signaling potency between type I and III IFNs is made more perplexing by the fact that both families share an identical intracellular Janus kinase/Signal Transducer and Activator of Transcription (JAK/STAT) signaling pathway^27,28^. Cytokines utilize the JAK/STAT pathway to transmit signals to the intracellular domains to initiate signaling but do so by using different combinations of signaling proteins and complexes. For type I and III IFNs, the same pairing of JAK1 and TYK2 kinases is utilized. However, a lack of structural information regarding full-length JAK proteins in natural complexes with full-length cytokine receptors has rendered some finer aspects of the JAK/STAT pathway inaccessible. It has yet to be addressed if and how the JAK kinase domains reorient during and after ligand stimulation. Heterodimerization of receptors induced via ligand binding again prompts additional questions^29^. Canonically, cell signaling is initiated when two JAKs bound to respective receptors are brought within a defined distance for transphosphorylation to occur. However, our current understanding does not make clear how the kinase domains of complex sharing JAKs are oriented with respect to each other, and if the relative orientation differs amongst distinct cytokine receptors. We hypothesize that these fundamental issues concerning the intracellular geometry of JAKs can be of much significance in decoding the differences in the functional activities of type I and III IFNs, a model system of cytokine signaling, as well as broader cytokine systems.

To determine if alterations to the geometric alignment of complex-sharing receptors could facilitate efficient transphosphorylation of proximal JAKs, we modulated the intracellular register of the IFNλR1-JAK1 axis and observed the effects on downstream JAK activation and subsequent signaling outputs. Alanine insertion mutagenesis was used to engineer mutant IFNλR1 receptors with precise twists in their intracellular registers that could realign the proximal JAKs within the heterodimeric signaling complex. Results indicate that the wild-type receptor contains a suboptimal alignment of transphosphorylating JAKs that is responsible for the low potency of type III IFNs relative to the type I IFNs. Functional potency of type III IFNs was found to be significantly enhanced by fine-tuning the receptor intracellular geometry. In this study, when we utilize a high-affinity ligand to stabilize the receptor complex and optimize the internal JAK geometry through receptor rotation, the type III IFN matches the type I IFN signaling in all measures of activity.

## Results

To interrogate how the geometry of proximal JAKs within a signaling complex affects type III IFN signaling, we subjected the transmembrane domain of IFNλR1 to alanine-insertion mutagenesis – an approach which has been previously utilized to alter the intracellular register of a cytokine receptor (Fig. 1, a)^30,31^. We engineered mutant IFNλR1 receptors with 1-4 alanine residues inserted after V242 within the alpha-helical transmembrane domain, with each added residue rotating the intracellular register by 109-degrees (Fig. 1, b). These mutated receptors, referred to here as λR1-1A, λR1-2A, λR1-3A, λR1-4A, were expressed stably in human embryonic kidney (HEK) 293 cells which are normally non-responsive to type III IFNs due to their very low expression levels of IFNλR1 but become responsive after being transduced to express exogenous IFNλR1^4,32^. N-terminal Flag tags were incorporated to enable accurate quantification of receptor expression levels, which were then used to normalize functional data (S. Fig. 1, a to e).

**Figure 1:**
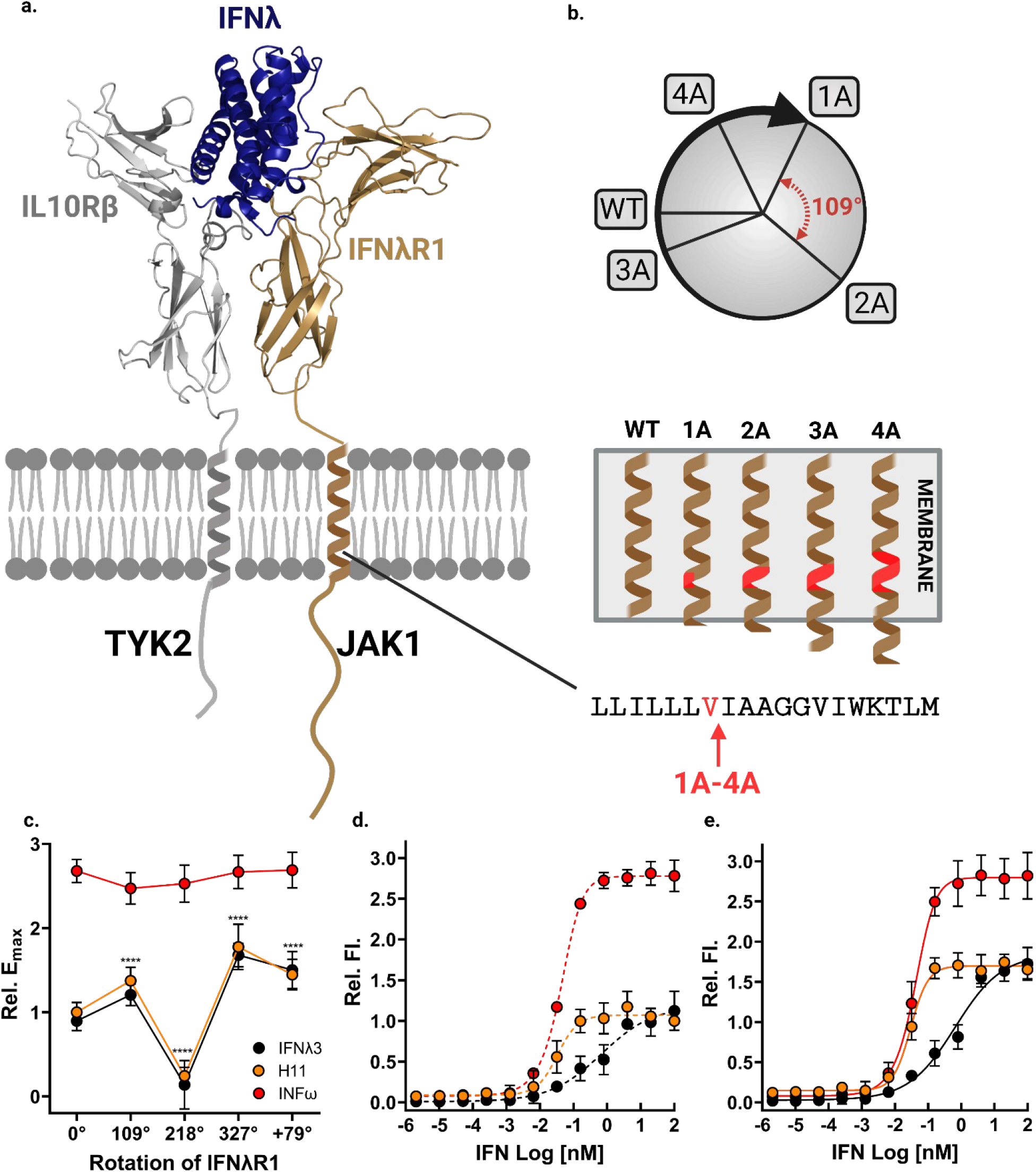
Modifications in the geometry of IFNλR1 modulate pSTAT1 responses. **a**, Schematic diagram of alanine insertion mutagenesis of the IFNλR1 transmembrane domain. **b**, α-helical wheel projections of the register rotations introduced by addition of each alanine residue are shown (top) and alanine residues (ranging from 1 to 4) were inserted after V242 (bottom). The direction of rotation is arbitrarily assigned with each residue adding a 109° rotation. **c**, Comparison of Emax values induced in the wild-type vs mutant IFNλR1 expressing cells by 1μM each of IFNω (red), IFNλ3 (black) or H11 (orange). All values were normalized to the wild-type receptor expressing cells treated with IFNλ3 (n=9). **d**, Relative quantification of pSTAT1 staining in cells expressing either the wild-type or **e**, mutant IFNλR1 with 3 alanine insertion by flow cytometry. Cells were treated with serial dilutions of IFNω (red), IFNλ3 (black) or H11 (orange) for 15 min. Curves were fit to a first-order logistic model. Error bars represent ± SEM (n=3). *p < 0.05; **p < 0.01; ***p < 0.001; ****p < 0.0001.

First we measured the direct downstream target of IFN signaling, phospho-STAT1 (pSTAT1) levels, in transduced HEK 293 cells treated with IFNω, wild-type IFNλ3 or engineered high-affinity IFNλ3 variant or H11 that was previously reported^33^. As predicted, the E_max_ values (maximum signaling potency) of pSTAT1 induced by IFNω were largely unaffected in all cell lines (Fig. 1, c). In contrast, cells treated with IFNλs differed in their maximal signaling amplitudes. IFNλR1 receptors with 2-alanines inserted in the transmembrane domain resulted in a near total loss of pSTAT1 signaling whereas cells with 3-alanines inserted resulted in the doubling of the E_max_ value compared to wild-type receptor expressing cells (Fig. 1, c). Insertion of either 1 or 4-alanines also significantly improved signaling, with λR1-4A cell line also outperforming the suboptimal signaling imposed by the JAK-JAK geometry in the wild-type receptor complex.

In accordance with existing literature, we found that type III IFNs trailed significantly behind type I IFNs in terms of pSTAT1 signaling potency. The experimental E_max_ of IFNω was ~2.67 fold over that of wild-type IFNλ3 in wild-type IFNλR1 expressing cells^33^. Notably, the fold difference is reduced to ~1.5 fold difference relative to the type I IFNs by having IFNλ3 signal through the mutant IFNλR1 with three added alanine residues. It should be noted that while the signaling maxima for both IFNλ3 and H11 ligands were markedly increased by the change in the receptor orientation, EC_50_ values remained largely unperturbed (Fig. 1, d and e). Together with our previous findings, these results indicate that the geometry of the intracellular receptor-JAK complex determines the strength of the signaling while the stability of the extracellular receptor-ligand complex, the cellular sensitivity^33^.

Based on the results, the data suggest that the relative positioning of intracellular JAKs in native heterodimeric complexes of type III IFNs results in non-optimal downstream signaling. We have shown here that we can effectively rotate the intracellular register of IFNλR1 by introducing helical twists in the transmembrane domain of the receptor. The consequences of the resultant rotations are particularly evident in cells expressing IFNλR1-2A (S. Fig. 2). In this cell line, pSTAT1 signaling is virtually lost due to a near 180-degree flip in the orientation of JAK1 with respect to TYK2, which likely poses a physical impossibility for the JAKs to transphosphorylate. On the contrary, when the register of IFNλR1 ICD is rotated by a predicted 327-degree from its native position, we observed a significant gain in the signaling amplitude. Taken collectively, we reason that for optimal type III IFN signaling, the complex-sharing JAKs must not only be within a defined distance but also that the proximity must be complemented with the proper register through which the cytokine signaling can be tuned.

Antiviral signaling assays were used to determine if the antiviral potency of type III IFNs are improved by tuning the signaling amplitude through the use of geometry-optimized IFNλR1 receptors. The wild-type and optimized IFNλR1 expressing cell lines were infected with a recombinant vesicular stomatitis virus linked to a green fluorescent protein construct (VSV-GFP) (Fig. 2, b). Consistent with the pSTAT1 signaling profiles, type III IFNs showed complete loss of antiviral activity in the IFNλR1-2A cells (S. Fig. 3) whereas IFNω induced similar antiviral responses across all cell lines (Fig. 2, c and f). Interestingly, the wild-type IFNλ3 and its high-affinity variant, H11, which were 84 and 12-fold lower in activity (EC_50_) than IFNω respectively in wild-type IFNλR1 expressing cells, effectively matched their antiviral activities to IFNω in IFNλR1-3A expressing cells (Fig. 2, a).

**Figure 2:**
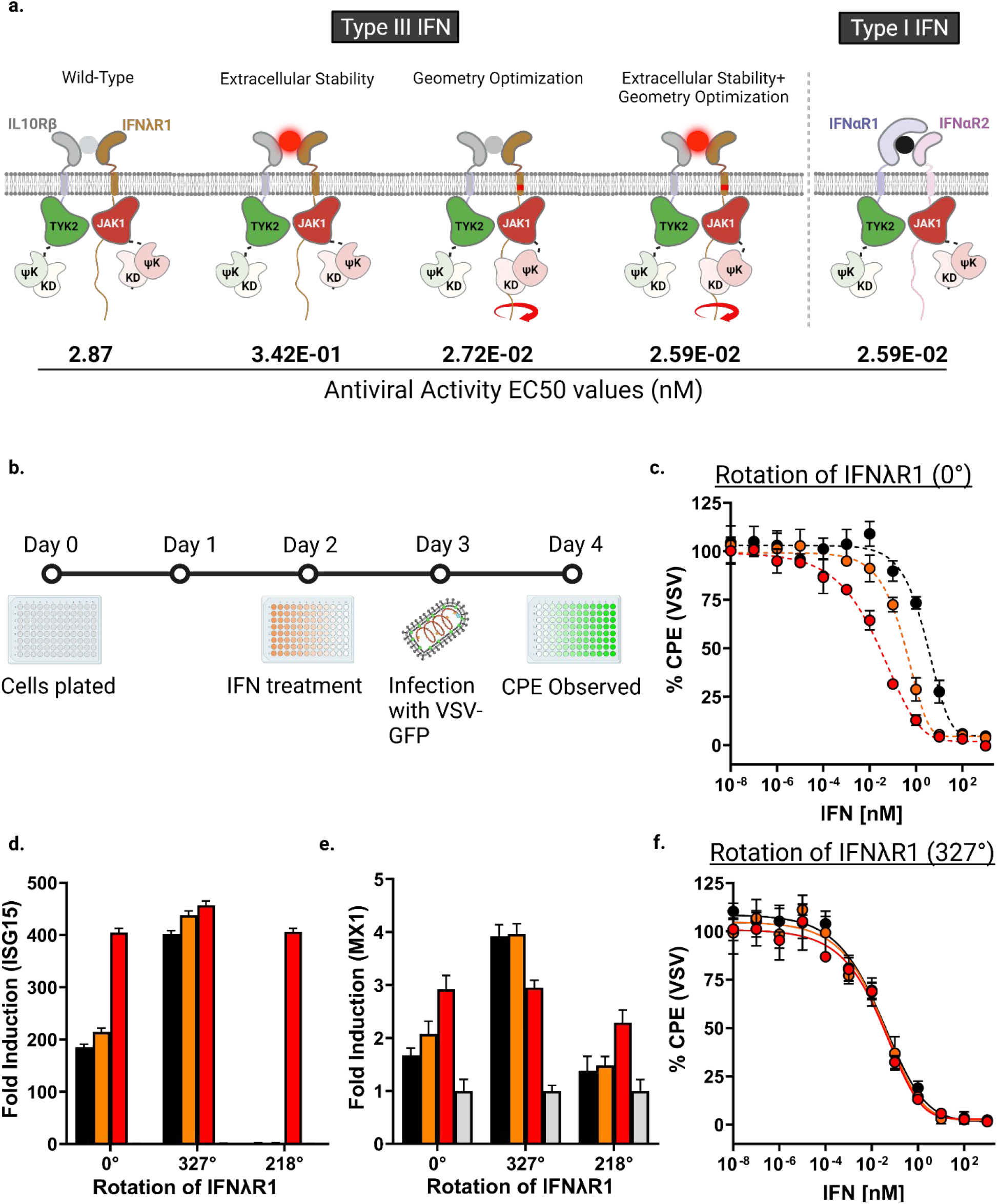
Register optimization improves type III IFN antiviral response against VSV infection. **a**, Schematic diagram summarizing the optimization strategies and their respectively associated EC50 values (nM) of the antiviral assay and calculated fold-changes relative to IFNω treated wild-type IFNλR1 expressing cells (assigned value 1). **b**, Schematic diagram depicting the antiviral assay set up. **c**, Antiviral activity of IFNs in cells expressing either the wild-type or **f**, optimized IFNλR1. Cells were incubated with serial dilutions of IFNλ3 (black), H11 (orange) or IFNω (red) for 24h prior to VSV-GFP viral infection at 80,000 PFU/well. Fluorescence levels were recorded 18h post-infection. Curves were fit to a first-order logistic model. Error bars represent ± SEM (n=3). **d**, PCR quantification of fold changes in induction of ISG15, **e**, MX1 at 6h post treatment with 100nM each of IFNλ3 (black), H11 (orange), or IFNω (red) in wild-type or optimized cell lines. Mean changes ± SEM in gene expression were determined relative to untreated cells (grey, assigned value of 1) and normalized to 18S (n=4).

A previous study conducted in HCV-infected Huh7.5 cells reported that the antiviral activities of type III IFNs can be improved 12-fold by a 150-fold improvement in the stability of the extracellular heterodimeric receptor complex through the engineered high-affinity ligand, IFNλ3 H11^33^. However, despite the significant gain in antiviral activity, the EC_50_ for IFNλ3 H11 remained 10-fold greater than IFNω^33^. Here, we show that the antiviral activity of type III IFNs is highly responsive to the increased pSTAT1 signaling potency modulated by the change in the intracellular IFNλR1 register. Notably, our results indicate that optimization of intracellular receptor-JAK geometry can potentiate the antiviral activities of type III IFNs, regardless of their receptor affinities, matching the activity achieved with type I IFNs.

In addition to their most prominent role as antivirals, type I IFNs are also known for their antitumor properties^34^. Here, we found that the anti-proliferative activities were most efficiently induced by IFNω across all cell lines (Fig. 3, c and f). The activity of IFNω is approximately ~8,500-fold (4-logs) over that of wild-type IFNλ3 signaling through wild-type IFNλR1. Analogous to pSTAT1 signaling, IFNλR1-2A expressing cells displayed no measurable anti-proliferative activity when treated with IFNλs (S. Fig. 4). Remarkably, the near 4-log difference in activities between the type I and III IFN was reduced to just ~30-fold in optimized IFNλR1 expressing cells stimulated with high-affinity H11 ligand, which represents a >280-fold, 3-log, improvement in activity (Fig. 3, a and f).

**Figure 3:**
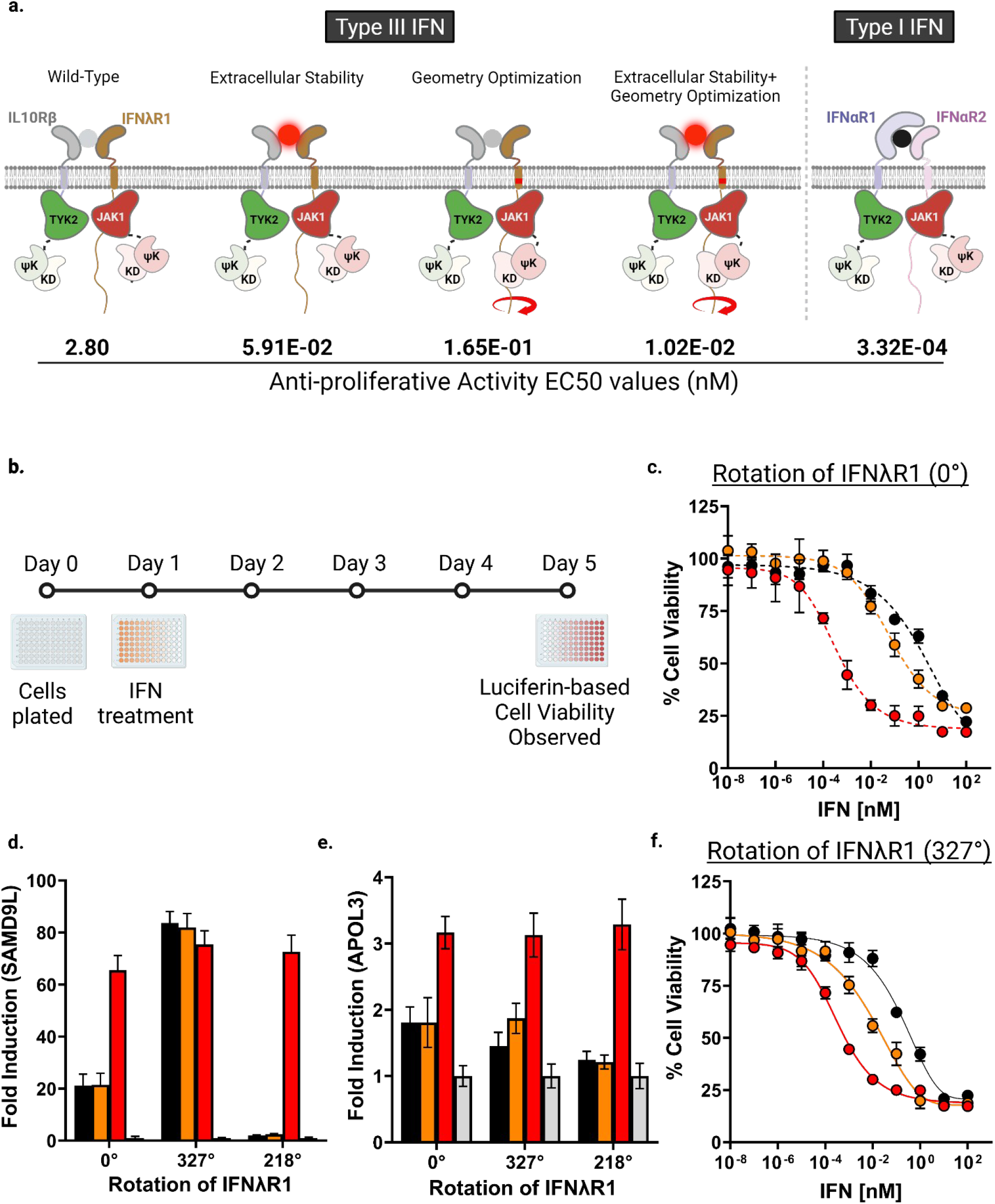
Optimization of IFNλR1 upregulates anti-proliferative activities of type III IFNs. **a**, Schematic diagram summarizing the optimization strategies and their respectively associated EC_50_ values (nM) of the anti-proliferative assay and calculated fold-changes relative to IFNω treated wild-type IFNλR1 expressing cells (assigned value 1). **b**, schematic diagram showing the experimental set-up of the assay. **c**, Anti-proliferative activity of IFNs in cells expressing either the wild-type or **f**, optimized IFNλR1. Cells were incubated with serial dilutions of IFNλ3 (black), H11 (orange) or IFNω (red) for 4 days. Curves were fit to a first-order logistic model. Error bars represent ± SEM (n=3). **d**, PCR quantification of fold changes in induction of SAMD9L, **e**, APOL3 at 6h post treatment with 100nM each of IFNλ3 (black), H11 (orange), or IFNω (red) in wild-type or optimized cell lines. Mean changes ± SEM in gene expression were determined relative to untreated cells (grey, assigned value of 1) and normalized to 18S (n=4).

It has previously been shown that the anti-proliferative activity of type III IFNs can be modulated via cell surface receptor overexpression and by enhancing the stability of the extracellular complex, with the former inviable as a method to potentiate signaling via adjuvants or other treatments ^32,34–36^. Here, our results indicate that the geometry of the intracellular signaling components majorly contributes to the anti-proliferative response and provides a viable means of therapeutic targeting. Furthermore, an analysis of the fold-changes in activity induced in different cell lines by two IFNλ ligands suggests a possible synergy between the affinity of ligand and the register of receptor-JAK complex in modulating anti-proliferative activities. When evaluated on the basis of receptor usage, the fold-increase in activity in response to the change in ligand affinity was significantly lower in the wild-type cells than in optimized IFNλR1 expressing cells. Similarly, when evaluated on the basis of ligand usage, the fold-increase in response to the change in receptor geometry was significantly lower for IFNλ3 than its high-affinity counterpart, H11. Maximal anti-proliferative effects were achieved only when the stimulation with high-affinity ligand was accompanied by cell signaling through geometry-optimized mutant IFNλR1 receptor.

We then quantified the transcriptional levels of a representative set of antiviral and anti-proliferative genes. At 6-hr post treatment with IFNs, the induction levels of antiviral genes (ISG15 and MX1), were significantly increased from 3 to 8-fold in optimized IFNλR1 expressing cells compared to the wild type (Fig. 2, d and e). Most notably, the gene induction levels by the high-affinity H11 effectively matched those of type I IFN, IFNω, in optimized IFNλR1 cells. Similarly, for anti-proliferative genes (APOL3 and SAMD9L), we observed fold-increases from 4 to 6-fold in gene induction levels by type III IFNs in the optimized IFNλR1 cells (Fig. 3, d and e). However, unlike the antiviral genes, the increased gene induction levels by type III IFNs, although greatly elevated, were only a fraction of those exhibited by IFNω. Notably, at 24-hr post IFN treatment, type III IFNs, regardless of their receptor affinity, induced all genes except APOL3 to near equivalent levels of type I IFN (S. Fig. 5, a to d). The gene induction study again highlights the importance of receptor geometry in downstream signaling outputs by IFNs. It should be noted that the two antiviral genes screened in this assay were more sensitive to the optimized register of the receptor than the anti-proliferative genes. This is consistent with our prior functional assays evaluating the antiviral and anti-proliferative activities of IFNs *in vitro*.

Next, we carried out next generation RNA sequencing to further evaluate the differences in the transcriptional responses to type I and III IFNs in wild-type vs optimized IFNλR1 expressing cells. Whole-genome transcriptional profiling of IFN treated cells after 24hr displayed strong correlation with the expression patterns of select antiviral and anti-proliferative gene sets obtained by qPCR. Principal component analysis (PCA) revealed four clusters – 1) untreated wild-type and optimized IFNλR1 cells, 2) IFNλ3 and H11 treated wild-type IFNλR1 cells, 3) IFNλ3 treated optimized IFNλR1 cells, and 4) IFNω treated wild-type and optimized IFNλR1 cells and H11 treated mutant IFNλR1 cells (Fig. 4, a and b). Compared to wild-type cells, optimized IFNλR1 cells displayed a significant increase in the abundance of overlapping differentially expressed genes (DEG) between type I and III IFNs treatments (Fig. 4, c). Overall, there was a significant increase in the number of upregulated genes in response to type III IFNs in optimized IFNλR1 cells than in wild-type cells (Fig. 4, d). Specifically, the number and fold-change of core antiviral ISGs were markedly improved for both IFNλ3 and H11 simulation in cells expressing optimized IFNλR1 receptors than the wild-type counterparts (Fig. 4, e to j).

**Figure 4.**
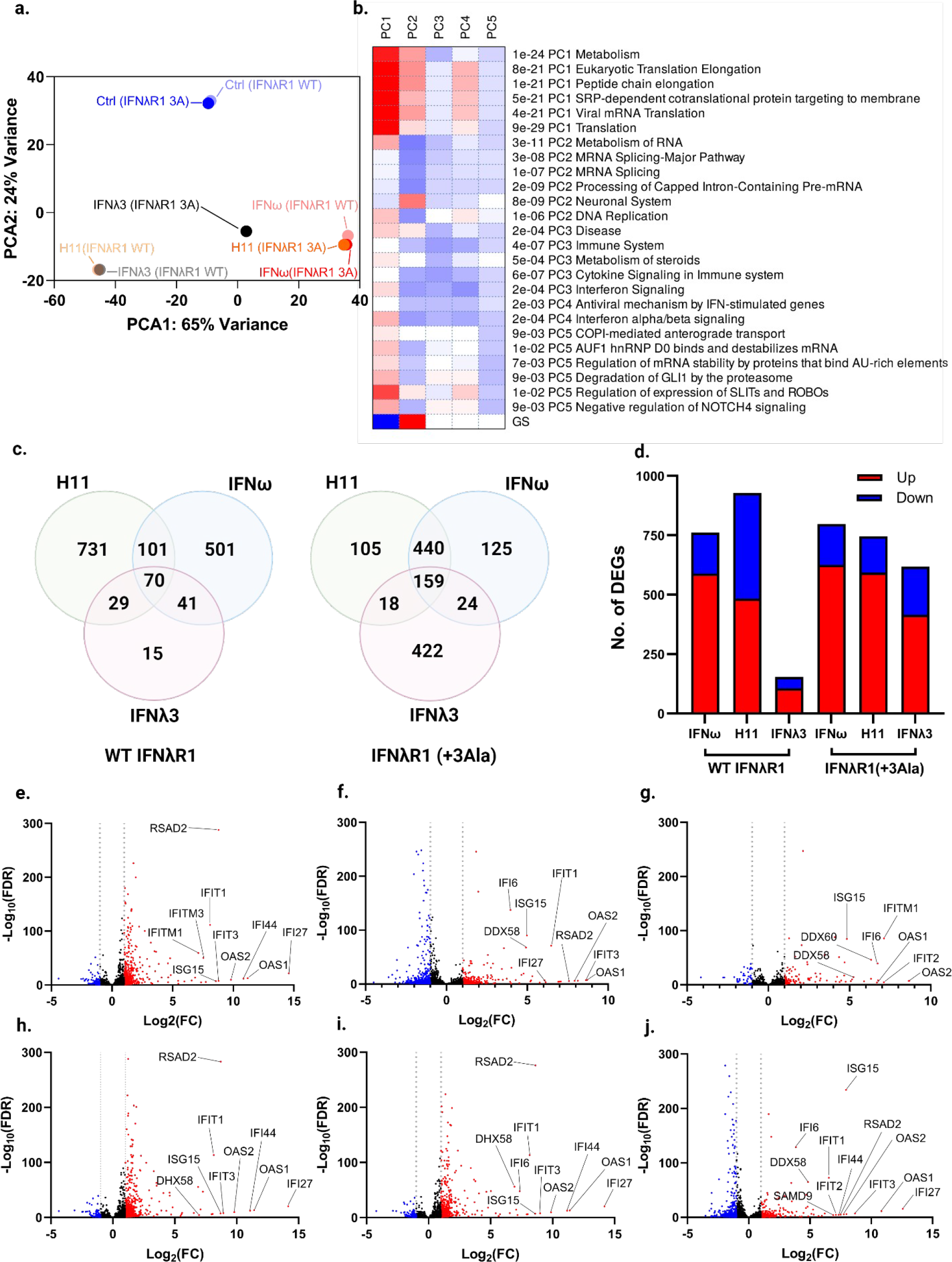
Transcriptome analysis indicates near type I IFN-level gene expression profile in optimized cells treated with high-affinity IFNλ3. **a**, PCA plot showing the distribution of wild-type and optimized IFNλR1 cell clusters treated with IFNω, IFNλ3 and H11 ligands for 24h. **b**, A heatmap showing the pathways involved in PCA analysis. **c**, Venn diagrams comparing the number of upregulated genes in cells expressing either wild-type (left) or optimized (right) IFNλR1 treated with indicated IFNs. **d**, A bar plot showing the quantification of DEGs in IFN-treated cells compared to untreated controls. **e-g**, Volcano plots showing decreased (blue) and increased (red) gene expression levels in cells expressing either wild-type or **h-j**, optimized (bottom) IFNλR1 treated with IFNω, H11 or IFNλ3 (left to right) compared to untreated controls. DE cutoffs were set at a log2 fold change of |1| and adjusted p value < 0.01.

K-means clustering analysis of 2,400 most variable genes indicated six distinct enriched pathways (S. Fig. 6, a). In both cell lines, type I IFN treatment led to the activation of genes involved in pathogen sensing and antigen processing/presentation (cluster I, black), innate and adaptive immune responses to viral infections (cluster II, green), and tissue repair and barrier functions (cluster III and IV, brown and yellow respectively) (S. Fig. 6, b to e). On the contrary, significant differences were observed for type III IFNs in the transcriptional profiles between wild-type and optimized IFNλR1 expressing cells. As anticipated, type III IFNs induced much weaker transcriptional programs of antiviral and barrier function-associated genes than type I IFN in wild-type cells. In optimized IFNλR1 expressing cells however, stimulation with H11 led to similar activation levels of gene subsets as type I IFN across all four clusters whereas stimulation with wild-type IFNλ3 showed a marked elevation in transcription levels compared to wild-type cells, although comparatively weaker than H11 or IFNω. Similarly, the activation states of core antiviral and anti-proliferative genes displayed clear enrichment patterns for IFNω and type III IFNs signaling through rotated IFNλR1 receptors (Fig. 5, a). Analysis of log2-transformed fold changes in individual select ISGs further indicated largely equivalent gene expression profiles among different IFN treatments in optimized IFNλR1 expressing cell lines (Fig. 5, b to g). Through Ingenuity Pathway Analysis (IPA), we further quantified the activation state of individual pathways involved in maintaining a state of immunity against pathogenic stimuli (Fig. 5, h). Consistent with our previous analyses, we observed an overlap in the enrichment of genes central to IFN-mediated antiviral responses between type I IFN treated cells and H11 treated optimized IFNλR1 expressing cells.

**Figure 5.**
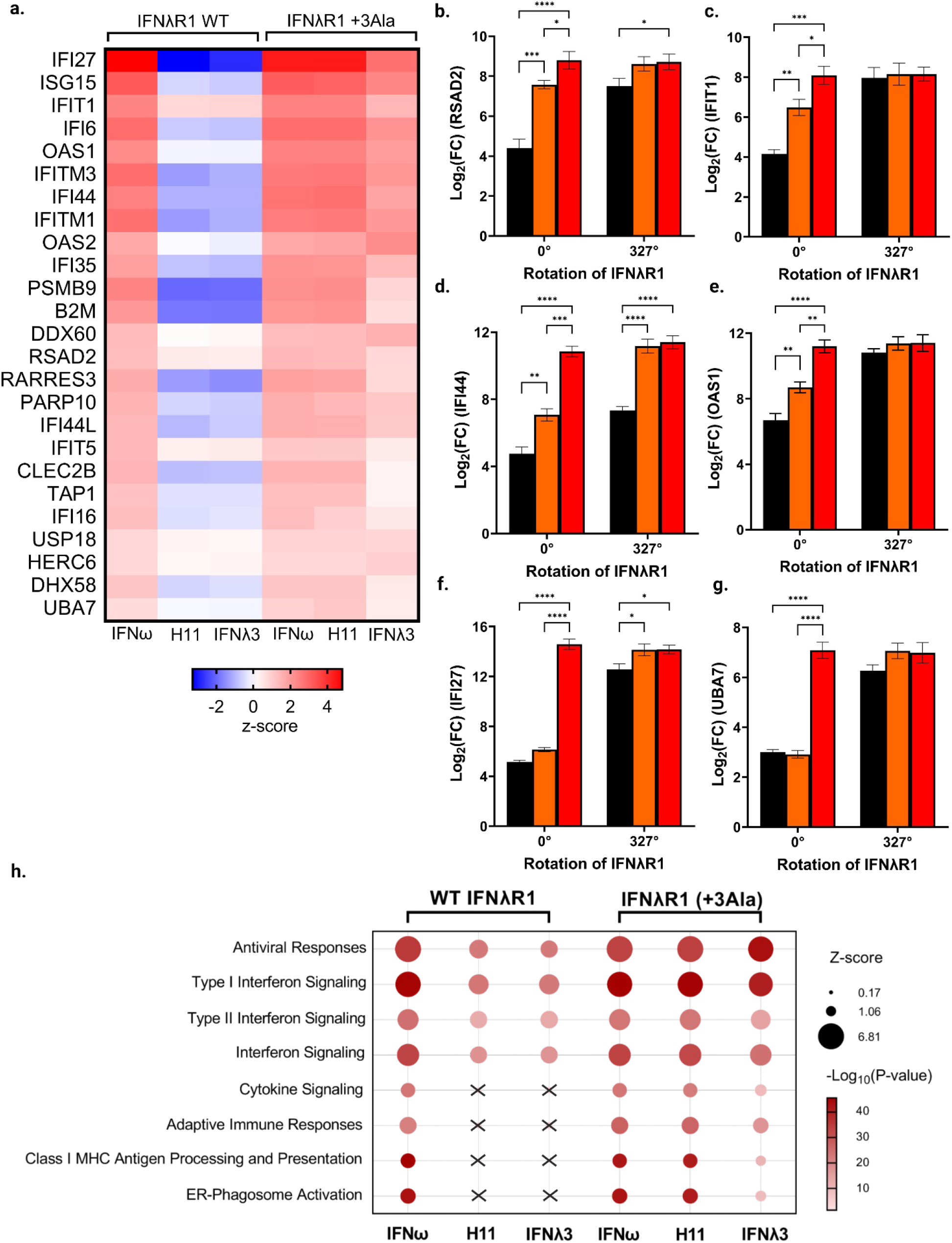
IPA analysis reveals similar ISG transcriptional profiles between type I and III IFNs in geometry optimized cells. **a**, Heatmap representation of activation levels of individual ISGs in cells expressing wild-type or optimized IFNλR1 treated with indicated IFNs. **b**, Log2-transformed relative expression of select antiviral ISGs including RSAD2, **c**, OAS1, **d**, IFT1 and anti-proliferative ISGS including **e**, IFI27, **f**, IFI44, and **g**, UBA7 in wild-type or mutant cell lines treated with IFNλ3 (black), H11 (orange), or IFNω (red). Statistical significance was determined by two-way ANOVA test. **h**, Bubble plot representation of significantly enriched antiviral mechanisms using IPA. Bubble color represents activation Z scores, and bubble size represents the −Log10 p value of enrichment. Statistical significance was determined by an activation Z score > |1| and a −log10 p > 1.32, which correspond to a p value of 0.05. Increases in −log10 p value are indicative of increased statistical significance.

**Figure 6:**
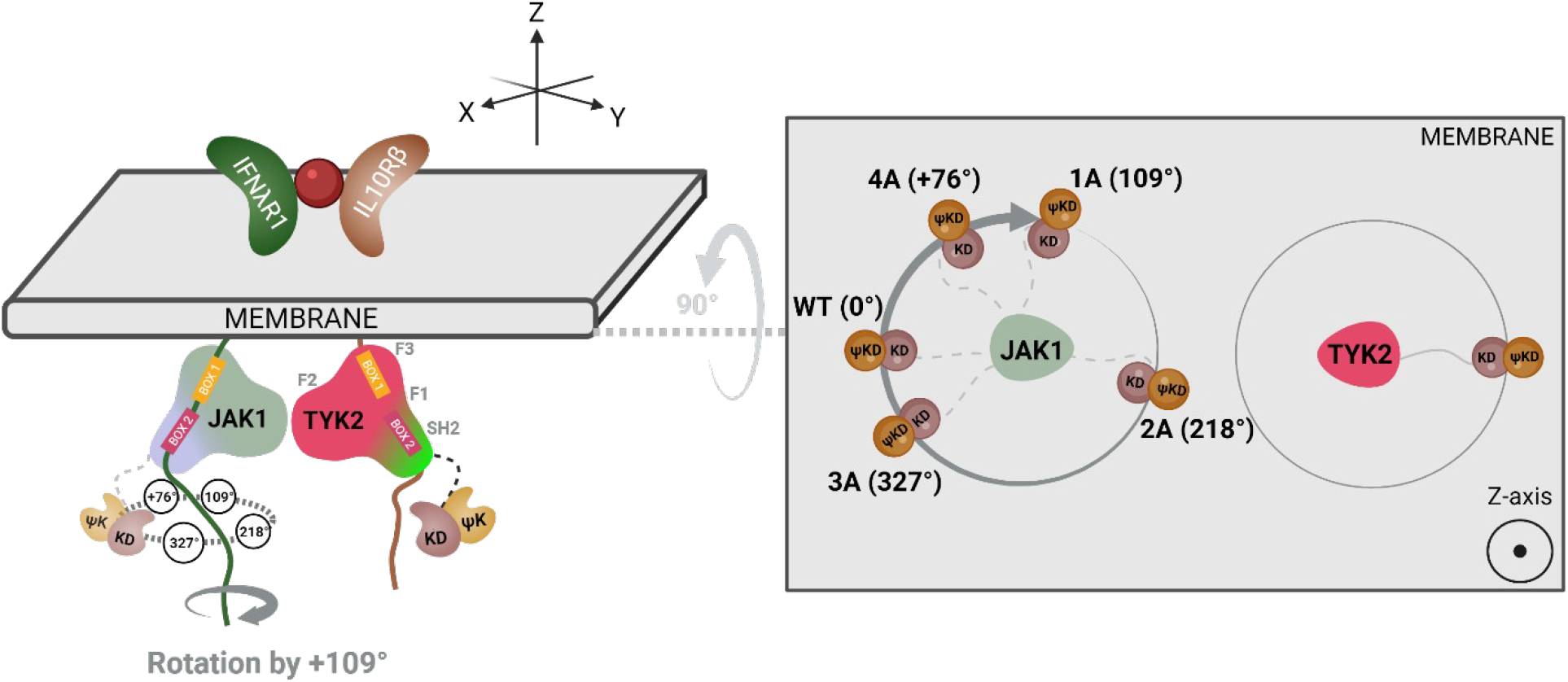
A schematic diagram shows the proposed positioning of C-terminal kinase domain of JAK1 relative to its N-terminal FERM SH2 domain when viewed down the axis of rotation. When 2 alanine residues are inserted in the transmembrane region of IFN-λR1 receptor, the near 180-degree rotation to the intracellularly associated JAK1 orients the kinase domains of JAK1 and TYK2 in a front-to-back manner, posing a physical barrier to transphosphorylation. On the contrary, the 327-degree rotation afforded by 3 alanine insertion decreases the distance between the kinase domains of JAK1 and TYK2 within the signaling complex, facilitating a more efficient transphosphorylation that leads to enhanced biological activities for type III IFNs.

## Discussion

The intrinsic discrepancy between type I and III IFN signaling strength is of particular interest because the intracellular signaling machinery employed by the two systems is nearly identical. Unlike receptor tyrosine kinases (RTKs), type I and III IFN receptors lack intrinsic tyrosine kinase domains in the cytoplasmic regions of their polypeptide chains and signal through the JAK/STAT pathway to propagate signals to the cytoplasmic components of the cascade^37^. There are four JAK kinases–JAK1, JAK2, JAK3 and TYK2 -and seven STAT proteins – STAT1, STAT2, STAT3, STAT4, STAT5a, STAT5b, STAT6^38,39^. In the canonical model, cytoplasmic tails of cytokine receptors are constitutively associated with a specific member of JAK protein via membrane-proximal binding sites, forming a complex that is functionally equivalent to RTKs^40–42^. The ligand binding event oligomerizes the receptors, bringing the receptor-JAK complexes into close proximity and allowing the JAKs to transphosphorylate each other. Activated JAKs in turn phosphorylate specific tyrosine/serine residues on the cytoplasmic regions of the associated cytokine receptor, creating docking sites for the SH2 domains of STAT proteins^29,43^. Specific STATs are then phosphorylated and released to allow formation of homo-or heterodimeric STAT complexes^44,45^. These complexes then subsequently translocate to the nucleus and bind to target sequences in the genome to initiate gene transcription^46^. Canonically, type I and III IFN receptors both utilize the JAK1/TYK2 kinases to form the STAT1/STAT2/IRF-9 (Interferon Regulatory Factor-9) signaling complex, known collectively as ISGF3 (Interferon-Stimulated Gene Factor 3) complex, which is an integral transcriptional regulator of core ISG genes^28^.

While the stability of the extracellular ligand-receptor complexes is a contributing factor to the downstream signaling outputs, it has been shown that engineered IFNλ ligands with higher receptor affinities exhibit limited effects on bridging the potency gap between type I and III IFNs^33^. In a previous study, a high affinity variant of IFNλ3 (termed H11) was engineered to increase the overall stability of the IFNλ ternary complex by 150-fold compared to the wild-type. Despite improvements in antiviral and anti-proliferative activities, the overall signaling potency and functional responses of H11 were significantly lower than those afforded by type I IFNs^33^. Although receptor abundance also seems to be a limiting factor for certain IFN activities, we speculate that there may be other characteristics inherent to receptor/JAK interactions that can further shed light on the differential functional capabilities of type I and III IFNs. We hypothesize that there exists an optimal geometrical alignment for the juxtaposed JAKs within the same heterodimeric IFN signaling complex that favors efficient transphosphorylation. We sought to address the question of whether type III IFN signaling can be made more potent by fine-tuning the geometry of their intracellular signaling complex. Hence, using alanine insertion mutagenesis approach, we engineered mutant IFNλR1 receptors with precise twists in their intracellular registers.

We conducted a series of functional assays probing the pSTAT1 signaling, antiviral, anti-proliferative activities and ISG gene induction levels by stimulating wild-type and mutant IFNλR1 receptor-expressing cell lines with type I IFN (IFNω) and type III IFNs -wild-type IFNλ3 and high-affinity H11 ligands. Our cytopathogenicity assay in cells infected with recombinant VSV-GFP virus shows that the antiviral efficacy of type I IFNs was effectively matched by type III IFNs signaling through optimized IFNλR1 receptors. This represents a 111-fold improvement in EC_50_ values of type III IFNs to native receptor-expressing cells. Similarly, the anti-proliferative activities of type III IFNs show significant improvements in response to the change in the receptor register. In cells expressing optimized IFNλR1 receptors, the EC_50_ of high-affinity IFNλ3, H11, was only 30-fold lower than that of IFNω, which is a drastic reduction of the near 4-log activity gap between IFNω and IFNλ3 in wild-type receptor-expressing cells.

Consistent with the increased pSTAT1 E_max_ levels and improved functional activities, the IFN-inducible gene expression is also improved in optimized IFNλR1 expressing cells. At 6h post-stimulation with IFNs, the two antiviral genes (MX1 and ISG15) showed the most drastic differences between the wild-type and optimized receptor-expressing cell lines. Most notably, the high-affinity IFNλ3 H11 ligand was able to match the antiviral gene induction levels exhibited by IFNω. Though significantly improved by receptor reorientation, the anti-proliferative gene expression levels (SAMD9L and APOL3) of IFNλs, however, remained lower than those of type I IFNs. Interestingly, the transcriptional profiles obtained after 24h post-stimulation showed that IFNλs, regardless of their receptor affinities, induced expression of the antiviral (MX1 and ISG15) genes and SAMD9L, an anti-proliferative related gene, to equal extents as IFNω. Whole-genome transcriptional analysis also showed that genes associated with mounting innate and adaptive immune responses against viral infections as well as maintaining tissue barrier functions were markedly upregulated in cells expressing optimized IFNλR1 receptors when treated with type III IFNs. The extent of overall gene activation by the high-affinity ligand, H11, was nearly identical to that of IFNω whereas that by wild-type IFNλ3 was weaker. However, when the analysis is limited to core antiviral ISG subsets, the wild-type IFNλ3 was able to achieve similar expression levels as H11 by the optimization of JAK-JAK geometry, which is consistent with our antiviral assay in which both IFNλ3 and H11 displayed similar EC_50_ values toward VSV-GFP infection. The results indicate that while geometry optimization can significantly expand the number and fold-changes of ISGs activated by type III IFNs, the optimization must be accompanied by an improved extracellular complex stability through use of high-affinity ligands in order to match the transcriptional profile achieved by type I IFNs.

Here we propose that in the context of type III IFN signaling complexes, the rotational repositioning of the receptor bound JAK1 afforded by the insertion of 1, 3 or 4 alanine residues in the transmembrane region of IFNλR1 reorients the kinase domain of JAK1 relative to that of TYK2 bound to IL10Rβ in such a way that the activation loops from both kinase domains are brought into either a more ideal orientation and or a closer proximity. In the absence of high-resolution structures of the type I and type III IFN receptor complexes that include the intracellular JAKs, the absolute rationale as to the extent with which the increased proximity and/or the optimized orientation of JAKs influenc the strength of cytokine signaling remains difficult to be determined definitively. Ultimately, our results indicate that the receptor-JAK interface adds another layer of complexity in understanding the functional differences between type I and III IFNs.

Severe acute respiratory syndrome coronavirus 2 (SARS-CoV-2) global outbreak has so far claimed over 5 million people worldwide^47^. The pandemic has precipitated tremendous research efforts to develop vaccines and antiviral therapeutics to curb risk of transmission, disease severity and mortality rates. Given the successful mainstream applications of type I IFNs in the treatment of chronic hepatitis B and C viruses (HBV, HCV), SARS-CoV-2 outbreak has instigated renewed interest in both type I and III IFNs^22–24,48^. Thus far, a landmark study of IFNβ1a alone or in tandem with remdesivir (NCT04492475) conducted by National Institute of Allergy and Infectious Diseases (NIAID) associated worse clinical outcomes due to severe adverse events with IFNβ1a treatment^49^. On the contrary, although adverse effects are notably lower with pegylated IFNλ1, phase 2 clinical trial data have so far been mixed largely due to the low therapeutic efficacy of IFNλs^21,25^. In addition to their most prominent role as antivirals, type I IFNs are also used as anti-tumor agents^34^. Both the recombinant and pegylated forms of certain type I IFNs, IFNα subtypes in particular, have been in clinics for some cancers such as melanoma, hairy-cell leukemia and Kaposi’s sarcoma^3^. However, due to the near ubiquitous expression of type I IFN receptors in tissues, the systemic administration of type I IFNs inevitably leads to off-target side effects. Given the overlapping gene expression profile between type I and III IFNs, type III IFNs with their limited receptor distribution and tissue abundance are increasingly regarded as more specific and less toxic alternatives to type I IFNs^50^. In regard to both antiviral and anticancer applications, it is evident that strategies to enhance the potency of IFNλs may have important clinical and public health implications in current and emerging epidemics as well as in our ongoing effort toward cancer therapies.

Although our work is focused on type I and III IFNs, the engineering tools, workflow, and resultant insights into the intracellular signaling machinery can be applied to evaluate and/or further our understanding of broader cytokine systems. We demonstrate that by fine-tuning the receptor-JAK interactions, we can significantly narrow, if not entirely eliminate in certain aspects, the expansive gap in potency between type I and III IFNs. Ultimately, our findings have opened up future venues for continued research that will have significant impact not only on expanding our canonical understanding of the JAK/STAT pathway in the context of IFN signaling but also on devising novel strategies for clinical translations of type III IFNs and other cytokines through the approach presented here.

## Materials and Methods

### Cell lines and cell culture

Authentication of cell lines used in this study is guaranteed by the sources. SF9 cells in Sf-900™ II SFM (Gibco), Hi5 cells in Express Five™ Medium (Gibco) and HEK 293 cells in FreeStyle™ 293 Expression Medium (Gibco) were purchased from Thermo Fisher and maintained in their respective recommended media. Sf-900™ II SFM and Express Five media were supplemented with 50μg/mL gentamicin and FreeStyle™ 293 Expression medium, with 10U/mL of penicillin/streptomycin. Lenti-X 293T cells were a gift from Dr. Jun Huang of the University of Chicago and cultured in DMEM +10% fetal bovine serum. All cell lines were checked for mycoplasma contamination prior to usage.

### Protein expression and purification

IFN-ω and IFN-λ3 were expressed and purified using baculovirus expression system, as described previously^51^. Briefly, Hi5 express cells were infected with a pre-titered amount of baculovirus and cultured at 28°C for 72h before being harvested for proteins. The high-affinity IFN-λ3 variant, H11, was expressed similarly in HEK 293 cells. All proteins contained C-terminal hexa-histidine tags and were isolated by Ni-NTA affinity chromatography and further purified by size exclusion chromatography on a Superdex 200 column (GE Healthcare, UK), equilibrated in 10 mM HEPES (pH 7.4) and 150 mM NaCl. Proteins were stored in buffer with 10% added glycerol.

### Generation of lentivirus transduced mutant cell lines

For the generation of lentiviral pseudoparticles, Lenti-X 293T cells were plated in 6-well plates at a density of 0.6 × 10^6^ cell/mL overnight. Next day, the cells were co-transfected with a plasmid encoding a cytokine receptor of interest, packaging and envelope plasmids at a fixed ratio of 0.75/0.5/0.26μg per well respectively. For each transfection, 4.5uL Fugene HD transfection reagent (Promega) was combined with 1.5μg total DNA in 100μL of Opti-MEM (GIBCO). Cells were incubated with the transfection media for 3 days with added fresh media on day 2 before the supernatants were collected, passed through a 0.45μm filter and stored at −80°C in 10% FBS supplemented media. 1mL of lentivirus containing supernatant was used to transduce 1 × 10^6^ target cells with fresh media being added to transduced cells every 2-3 days. On day 5, stable expression of target receptors was determined by staining against their N-terminal Flag-tag with mouse anti-Flag conjugated to Alexa 488 (Abcam).

### *In vitro* pSTAT1 signaling assay

Cells were plated overnight in a 96-well format at a density of 10,000 cells/well and treated with serial dilutions of IFN-ω, wild-type IFN-λ3 or its high-affinity variant (H11) for 15 min at 37°C. The medium was removed, and cells were detached with Trypsin (Gibco) for 5 min at 37°C. Cells were transferred to a deep-well 96-well block containing an equal volume of 4% (w/v) paraformaldehyde (PFA) solution and incubated for 15 min at room temperature. Fixed cells were then washed three times with phosphate-buffered saline containing 0.5% (w/v) BSA (PBSA), resuspended in 100% methanol for 1h on ice. Cells were next stained with Alexa 488 conjugated pSTAT1 antibody (Cell Signaling Technology). The half-maximal response concentration (EC_50_) and E_max_ of signaling was determined by fitting the data to a sigmoidal dose–response curve (GraphPad Prism v.9).

### *In vitro* antiviral assay

Recombinant VSV harboring a green fluorescent protein (GFP) transgene (VSV-GFP) was a gift from Horvath lab, Northwestern University. HEK 293 cells were seeded at a density of 12,500 cells/well in a 96 well format and after 45h, the cells were treated with serial dilutions of IFN-ω, wild-type IFN-λ3 or its high-affinity variant (H11). Cell medium containing IFN treatment was removed after 24h and VSV-GFP virus diluted in serum-free media was added to the cells at 80,000 PFU/well. At 18h post-VSV-GFP infection, the cytopathic effects (CPE) were measured via a fluorescence plate reader.

### *In vitro* anti-proliferative assay

Cells were plated overnight in a 96-well format at a density of 10,000 cells/well. On the following day, the media was replaced with fresh media containing serial dilutions of IFN-ω, wild-type IFN-λ3 or its high-affinity variant (H11). Four days post IFN-treatment, cell density was measured using CellTiter-Glo (Promega) according to the manufacturer’s protocol.

### Quantification of gene induction by RT-qPCR

For measuring gene induction, 600,000 cells were plated in a 6-well format overnight and treated with 100nM each of IFN-ω, wild-type IFN-λ3 or IFN-λ3 H11 for 6 or 24h on the following day. RNA was extracted with the Monarch Total RNA miniprep kit T2010 (NEB), 1ug of which was converted to cDNA by a RT-PCR reaction using the High Capacity RNA-to-cDNA kit (Applied Biosystems). ISG induction relative to the untreated controls in wild-type cells was measured by qPCR assay (PowerSYBR Green PCR Master Mix, Applied Biosystems) on a QuantStudio 3 instrument (Thermo Fisher Scientific) following manufacturer’s instructions. Transcription quantification was normalized to 18S internal controls. Primers were purchased from Sigma-Aldrich. Please refer to the supplemental information Table 1 for a complete list of primers used.

### RNA sequencing and Transcriptome Analysis

Whole human transcriptome sequencing over 20,000 genes was performed on the Ion GeneStudio S5 Plus System using the Ion Ampliseq™ Transcriptome Gene Expression Kit (Thermo Fisher). Transcriptome libraries were barcoded, templated and sequenced using either Ion 550™ Kit-Chef and Ion 550 Chip Kit as one 16-plex library pool or Ion 540™ Kit-Chef and Ion 540 Chip Kit as one 8-plex library pool (Thermo Fisher). Two independent sequencing analyses were performed on a panel of eight samples. The RNA samples included in each panel are extracted from the following categories – untreated WT IFN-λR1 cells, untreated IFN-λR1 3A cells, IFNω treated WT IFN-λR1 cells, IFNω treated IFN-λR1 3A cells, IFN-λ3 treated WT IFN-λR1 cells, IFN-λ3 treated IFN-λR1 3A cells, H11 treated WT IFN-λR1 cells and H11 treated IFN-λR1 3A cells. Gene mapping and analysis was performed using Ion Torrent Suite™ v.5.10.0 (Thermo Fisher). Heat maps and figures showing PCA of gene expression were generated in MATLAB v.R2018b (MathWorks).

### Statistical Analyses

The results were presented as means ± standard deviation (STD). The statistical significance of differences between the groups was determined by two-way ANOVA analysis with subsequent correction for multiple comparisons using Tukey test. All statistical analyses were performed using GraphPad 9.0.2. Differences were considered statistically significant at \*\*\*\**p* < 0.0001, \*\*\**p* < 0.001, \*\**p* < 0.01 and * *p* < 0.05. The statistical analysis of experiments with technical replicates is detailed in figures’ legends.

## Data availability

The authors declare that all the data supporting the findings of this study are available within the article and its Supplementary Information files or from the corresponding author upon reasonable request.

## Acknowledgements

We thank Patrick Partisien from Horvath lab for providing the VSV-GFP virus stocks and an overview of the antiviral assay. We would also like to thank the University of Chicago and a grant from NIH NIGMS 1R35GM147179-01 for funding.

## Author contributions

T.A. and J.M. conceived the project. T.A. performed the experiments and analyzed the results.

T.A. and J.M. wrote the manuscript.

