## Supplemental for "Enhanced Complex Stability and Optimal JAK Geometry are Pivotal for a Potent Type III Interferon Response"

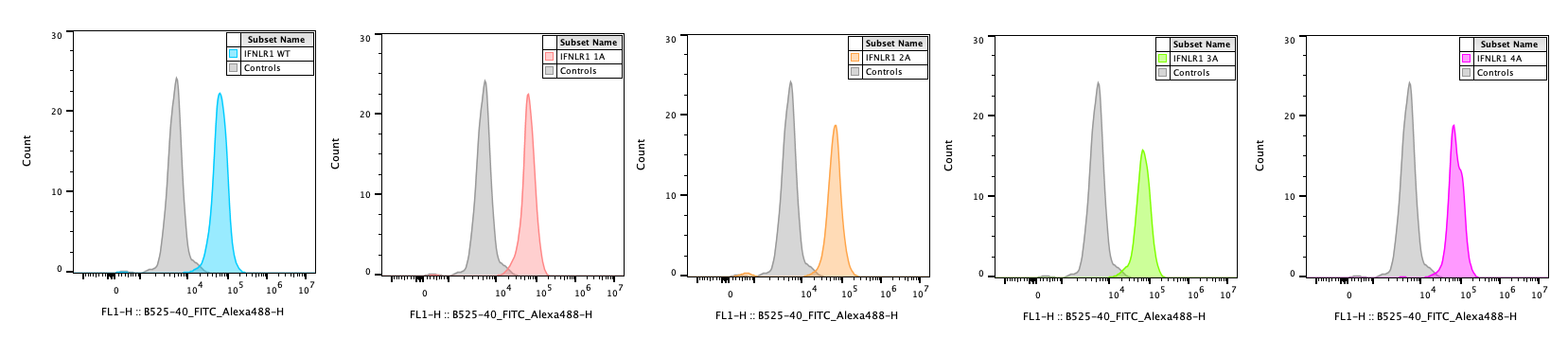

**Supplemental Figure 1**: Histograms depicting the surface levels of the wild-type or 1-4 alanine-inserted mutant IFNλR1 receptors (left to right) relative to non-transduced controls (grey). HEK 293 cells were transduced to stably express N-terminal Flag-tagged wild-type or mutant IFN-λR1 receptors, which were stained with anti-Flag conjugated to Alexa 488 and analyzed by flow cytometry.

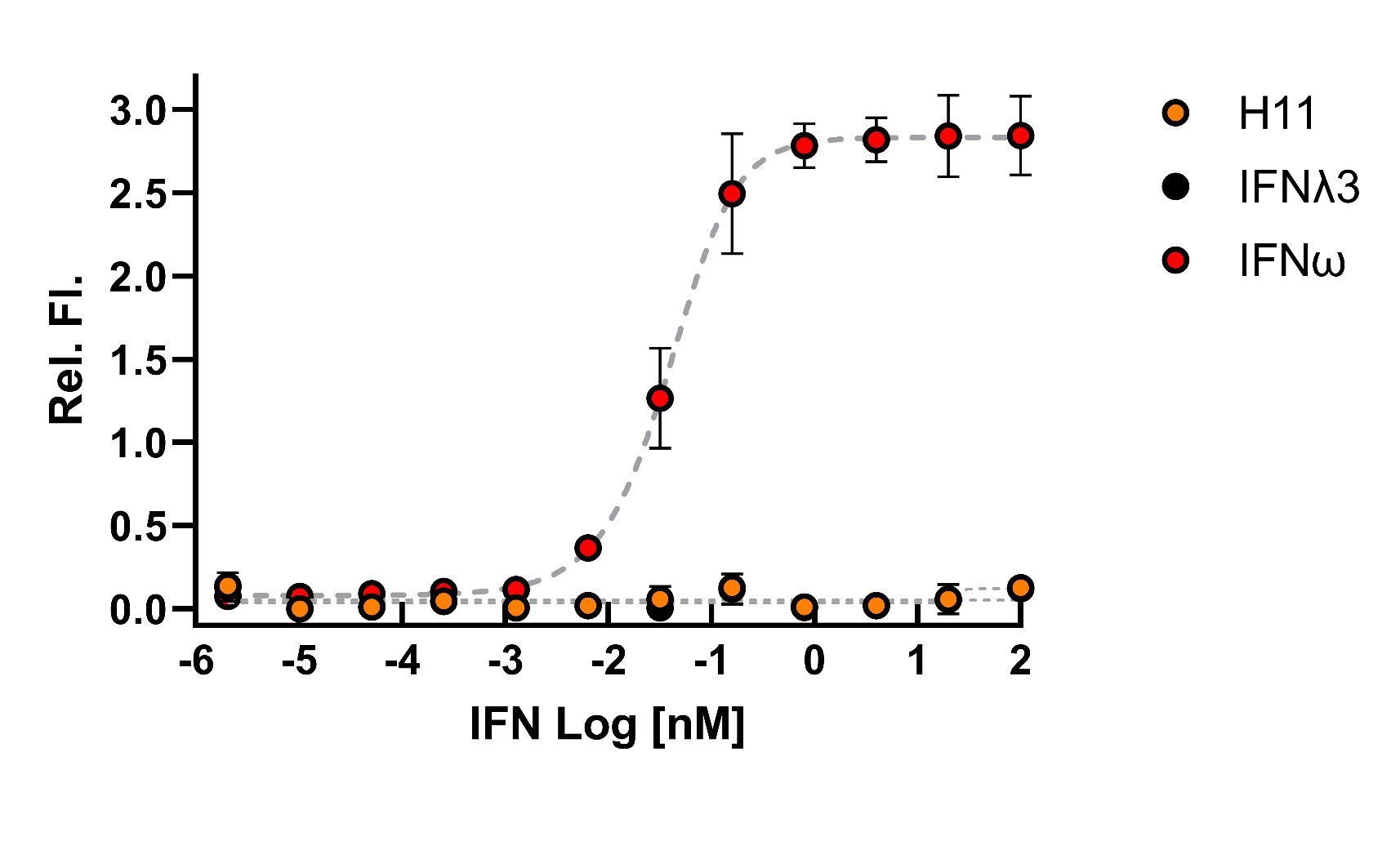

**Supplemental Figure 2:** Relative quantification of pSTAT1 staining in cells expressing mutant IFNλR1with 2 alanine insertion by flow cytometry. Cells were treated with serial dilutions of IFNω (red), IFNλ3 (black) or H11 (orange) for 15 min. Curves were fit to a first-order logistic model. Error bars represent ± SEM (n=3).

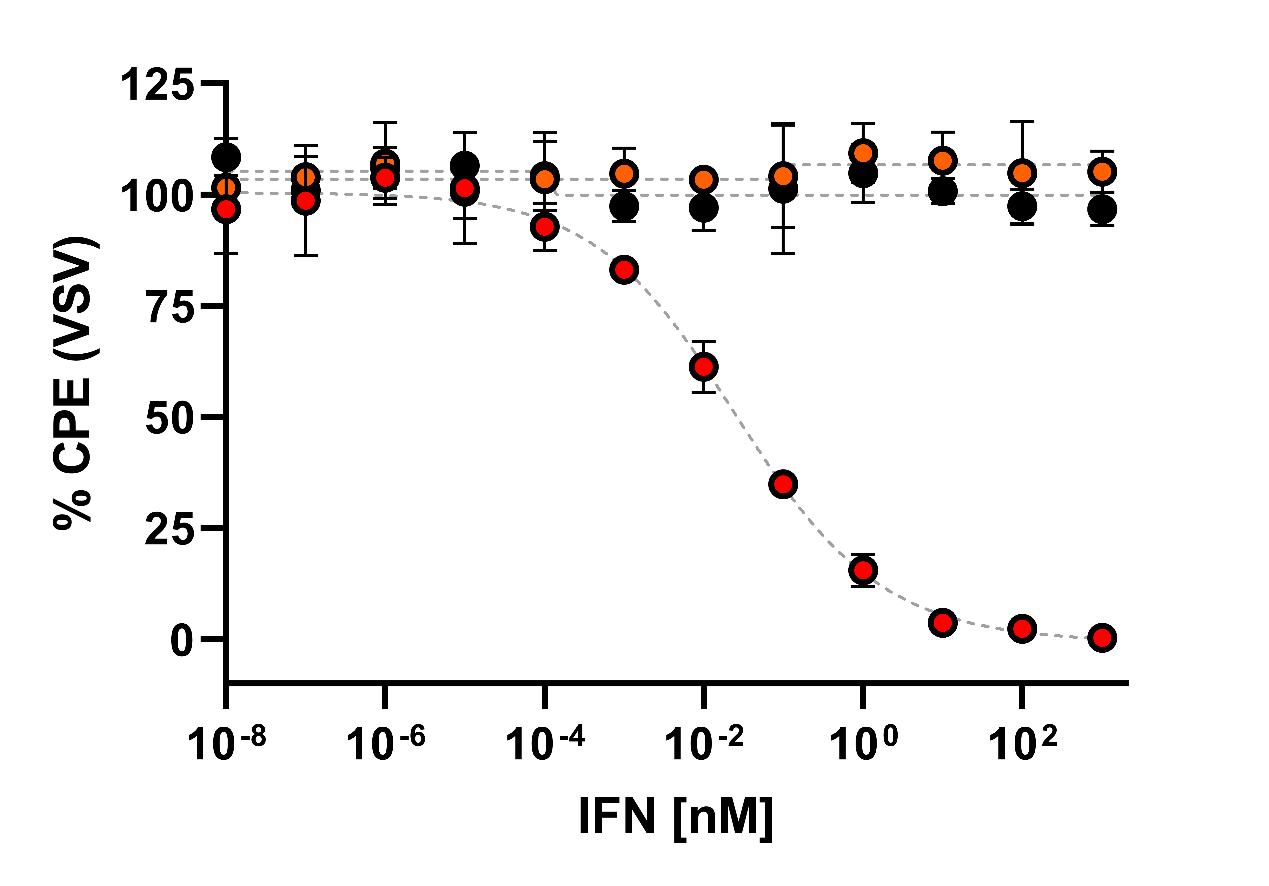

**Supplemental Figure 3:** Anti-viral activity of IFNs in cells expressing mutant IFNλR1 with 2 alanine insertion. Cells were incubated with serial dilutions of IFNλ3 (black), H11 (orange) or IFNω (red) for 4 days. Curves were fit to a first-order logistic model. Error bars represent ± SEM (n=3).

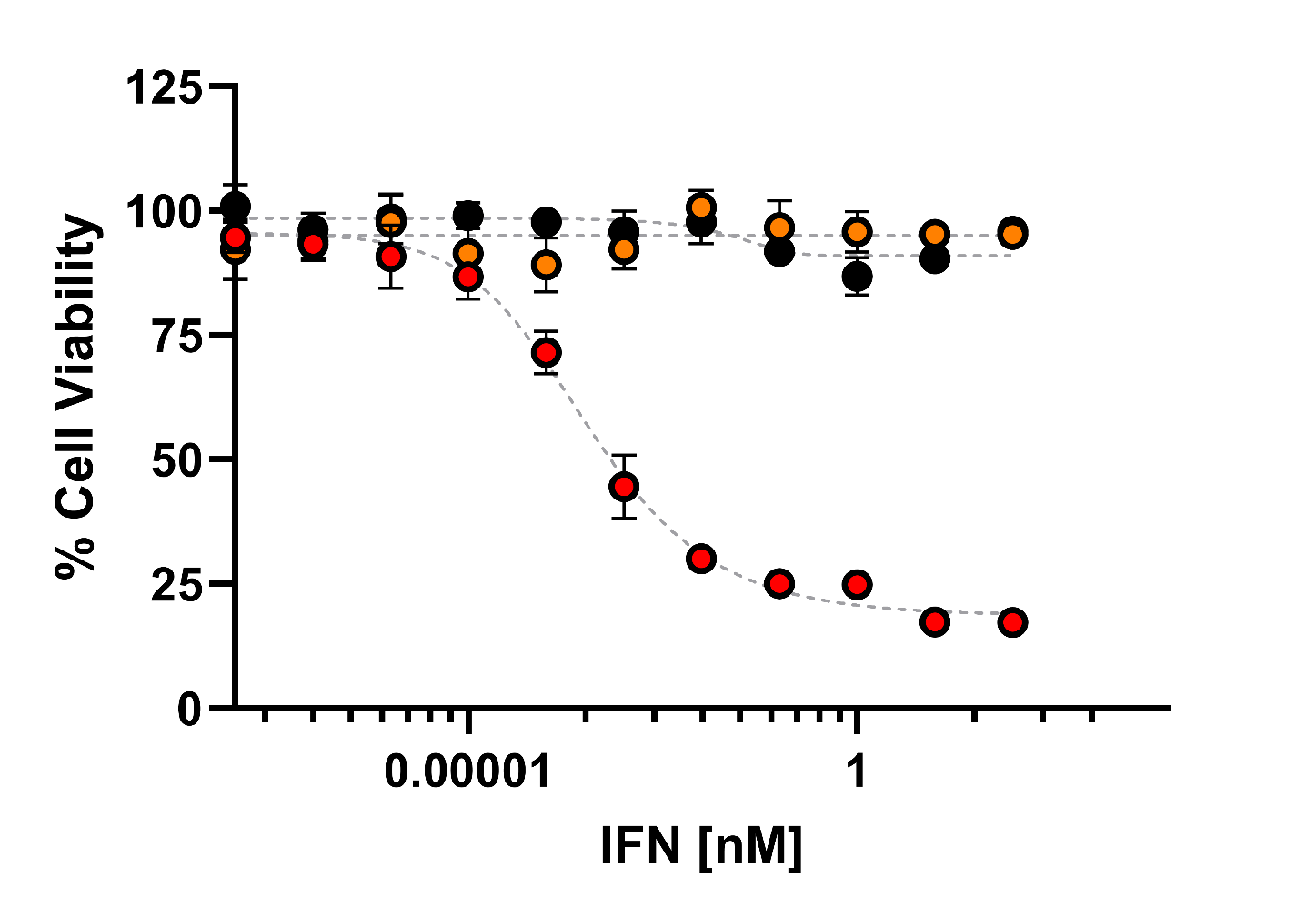

**Supplemental Figure 4:** Anti-proliferative activity of IFNs in cells expressing mutant IFNλR1 with 2 alanine insertion. Cells were incubated with serial dilutions of IFNλ3 (black), H11 (orange) or IFNω (red) for 4 days. Curves were fit to a first-order logistic model. Error bars represent ± SEM (n=3).

**c.**

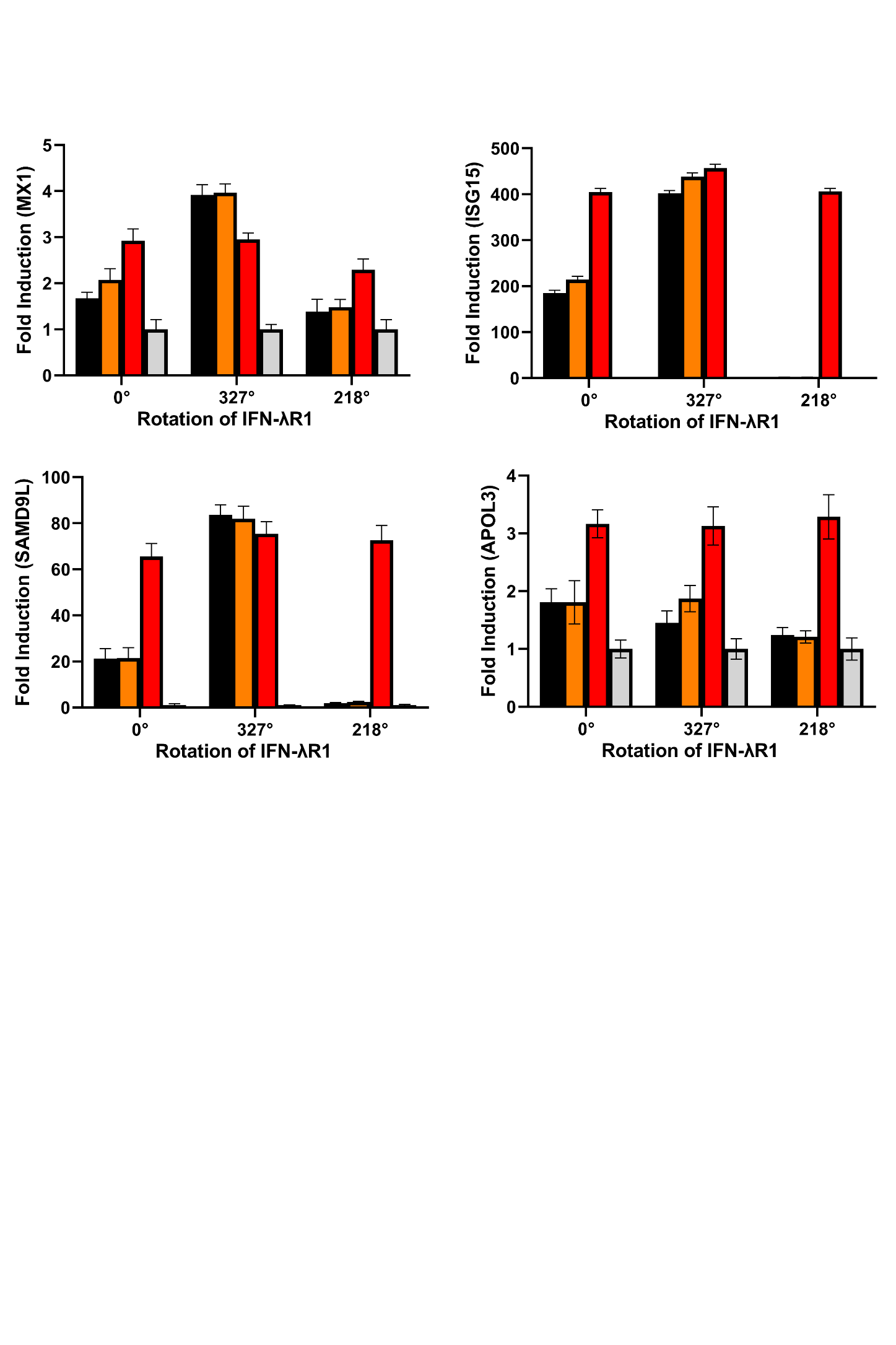

**a.**

**b.**

**d.**

**Supplemental Figure 5: a,** PCR quantification of fold changes in induction of MX1, **b**, ISG15, **c**, SAMD9L, **d**, APOL3 at 24h post treatment with 100nM each of IFNλ3 (black), H11 (orange), or IFNω (red) in wild-type or mutant cell lines. Mean changes ± SEM in gene expression were determined relative to untreated cells (grey, assigned value of 1) and normalized to 18S (n=4).

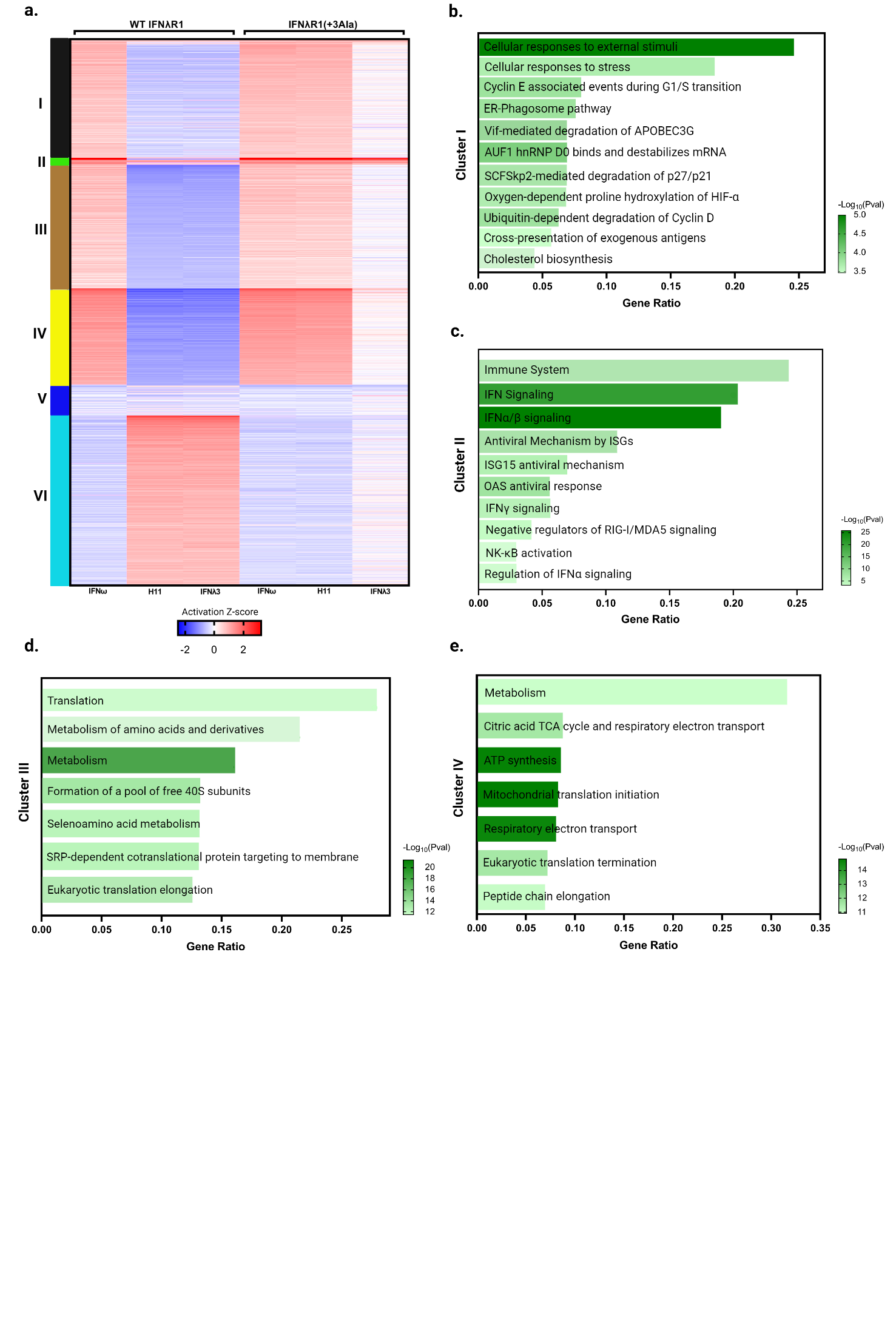

**Supplemental Figure 6: a,** Heatmap obtained by K-means clustering of 2400 genes showing six clusters. **b-e**, Bar plots detailing the enrichment pathways in the curated clusters. Bar size represents gene ratios within each enriched pathway, and color represents the -Log10 p value of enrichment. Increases in −log10 p value are indicative of increased statistical significance.

**a.**

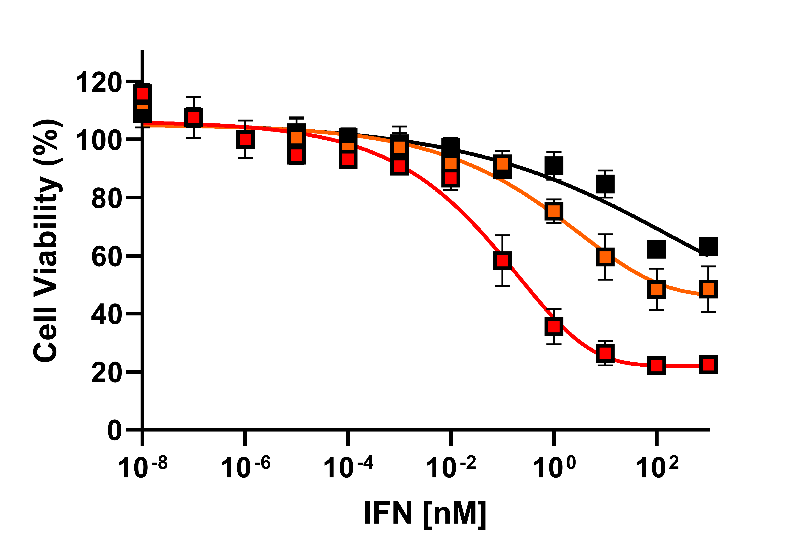

**b.**

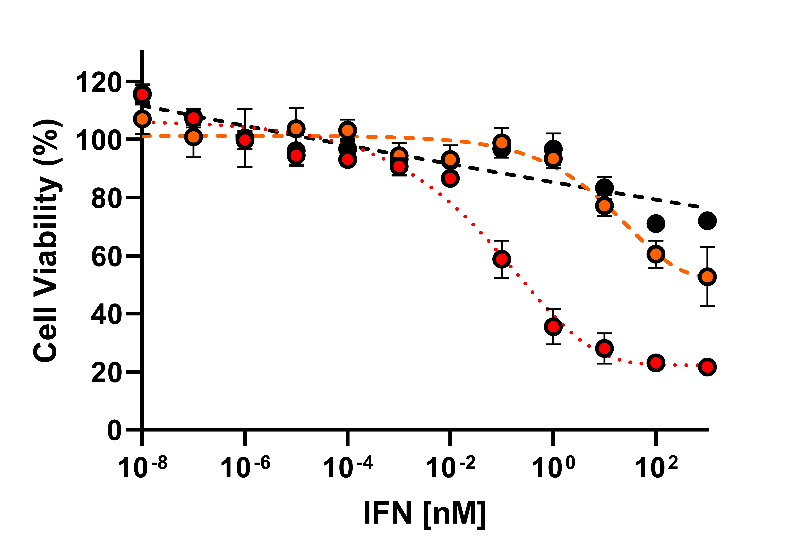

**Supplemental Figure 7:** **a**, Anti-proliferative activity of IFNs in MCF-7 cells expressing optimized or **b**, wild-type IFNλR1 receptors. Cells were incubated with serial dilutions of IFNλ3 (black), H11 (orange) or IFNω (red) for 4 days. Curves were fit to a first-order logistic model. Error bars represent ± SEM (n=3).
